# State-specific individualized functional networks form a predictive signature of brain state

**DOI:** 10.1101/372110

**Authors:** Mehraveh Salehi, Amin Karbasi, Daniel S. Barron, Dustin Scheinost, R. Todd Constable

## Abstract

There is extensive evidence that human brain functional organization is dynamic, varying within a subject as the brain switches between tasks demands. This functional organization also varies across subjects, even when they are all engaged in similar tasks. Currently, we lack a comprehensive model that unifies the two dimensions of variation (brain state and subject). Using fMRI data obtained across multiple task-evoked and rest conditions (which we operationally define as brain states) and across multiple subjects, we develop a state-and subject-specific functional network parcellation (the assignment of nodes to networks). Our parcellation approach provides a measure of how node-to-network assignment (NNA) changes across states and across subjects. We demonstrate that the brain’s functional networks are not spatially fixed, but reconfigure with brain state. This reconfiguration is robust and reliable to such an extent that it can be used to predict brain state with accuracies up to 97%.

## Introduction

The human brain is organized into functional networks that represent coordinated effort of individual subunits (or nodes) to execute specific functions (6-9). Previous studies have delineated 5-20 networks during the resting-state that putatively represent the “intrinsic” functional organization of the brain (9-12). These networks have been associated with a wide range of cognitive tasks (13-15) and alterations in the spatial organization of these networks have been linked to clinical disorders (16-18). Increasingly, there is evidence that a network’s spatial organization changes with the brain’s task demands (or state) (19-21). Characterizing a network’s state-evoked spatial reconfiguration is a key step towards fully understanding the brain’s functional organization. It is also essential to test the hypothesis that brain networks are not fixed but reconfigure as a function of brain state.

Recent studies have made significant progress in characterizing state-evoked changes in functional connectivity elicited by task performance (20-28). However, none of these works explicitly examined the possibility that networks spatially reconfigure: (i) The majority of these studies explicitly assumed that networks remain spatially unchanged across tasks (21, 22, 29). These analyses were restricted to investigating differences in connectivity between networks, and not whether networks spatially reconfigure across tasks. (ii) Another line of research takes a more abstract perspective by computing global network measures (such as modularity and participation coefficient) and comparing these measures across different states (27, 28, 30-32). Such approaches address the modular reconfiguration of the brain as a whole by defining a new set of networks for each state, but they do not quantify how or whether the same networks change across states. Such studies also typically define a small number of networks (~3-7) whose correspondence to the putative resting-state networks are unclear. These approaches cannot answer how specific networks reorganize as a function of brain state. (iii) Finally, many studies do not directly examine cross-subject variations in network reorganization (20), or treat cross-subject and cross-session changes as similar notions for defining reorganization (33). However, functional organization varies significantly across subjects, yielding large individual differences in network definitions (34, 35). There is a clear need to account for both cross-subject and cross-state variability in functional network organization when characterizing brain dynamics.

Here, we use a novel approach to dynamically map the brain’s functional networks across task-evoked and resting states and across individuals. We operationalize the brain’s functional subunits as a set of 268 pre-defined nodes (6). We then apply our recently developed exemplar-based parcellation method (36) to assign a set of pre-defined nodes to individualized, state-specific functional networks, while preserving the precise network correspondences across subjects and states. We show that all putative resting-state networks reconfigure by recruiting or dismissing nodes in a task specific manner as opposed to simply changing within and between network connectivity. We demonstrate the reliability of the observed state-evoked reconfiguration of networks with a fully cross-validated predictive model that decodes the brain state of unseen subjects solely based on their node-to-network assignments (NNA). We then divide nodes into three classes based on the stability of these NNAs: 1) steady nodes that exhibit the same network assignment across states and subjects; 2) flexible nodes that change their network assignment across states, but are consistent across subjects, and 3) transient nodes that change their network assignment across both subjects and states. To give behavioral context for these node classes, we use a large-scale meta-analysis of task activation studies from BrainMap to assign behavioral domains and paradigm classes to each node. Together, our findings provide a more comprehensive view of the dynamics of large-scale functional networks.

## Results

### Functional network configuration varies across states

We analyzed fMRI data from the Human Connectome Project (HCP; *n* = 718); each subject performed 2 rest (REST1 and REST2) and 7 task (MOTOR, GAMBLING, WORKING MEMORY [WM], EMOTION, LANGUAGE, RELATIONAL, and SOCIAL) runs (hereafter, “states”) in the scanner. We operationalized the brain’s subunits into 268 nodes using a whole-brain functional atlas defined previously in a separate sample (37). We calculated and normalized the squared Euclidean distances between the mean time courses of each node pair, generating nine functional distance matrices per subject. We used exemplar-based parcellation (36) to assign these nodes into functional networks in an individualized manner for each state and subject, such that every subject acquired an individualized node-to-network assignment (NNA) for each state. A significant advantage of this approach is that it allows for individualized networks while maintaining correspondence of networks across subjects and states. As the number of networks (K) is an arbitrary parameter, we repeated the analysis for K=2-25 networks. Results presented in the main text are for K=12 networks with results from different numbers of K networks presented as SI.

Figure 1a displays population-level, state-specific networks for K=12 networks across 9 states, computed using winner-takes-all strategy on all the subjects within each state. While a consistent network structure is apparent across states, clear differences in network membership are observed across all states and networks. For example, the posterior areas of the caudate nucleus are associated with the ventral attention network (VAN) during rest, but their network assignments change to the language or the sensorimotor network (SMN) during task performance (Figure 1a). To quantify network reconfiguration across states, we calculated the Hamming distance (38)—or the number of nodes that change their NNAs—for each pair of states (Figure 1b). The two resting-states (REST1 and REST2) have the most similar functional organization with only a 5.6% (15/268) difference in their NNAs. Given that the resting-state scans represent the same state, this result provides a benchmark for comparisons between different states. In contrast, the smallest reconfiguration between two different states was over three times larger: WM and GAMBLING, 19% (51/268). GAMBLING and SOCIAL demonstrate the highest level of network reconfiguration (45.5% difference). SOCIAL showed the least similarity to the other states with 41.9% difference on average.

**Figure 1.**
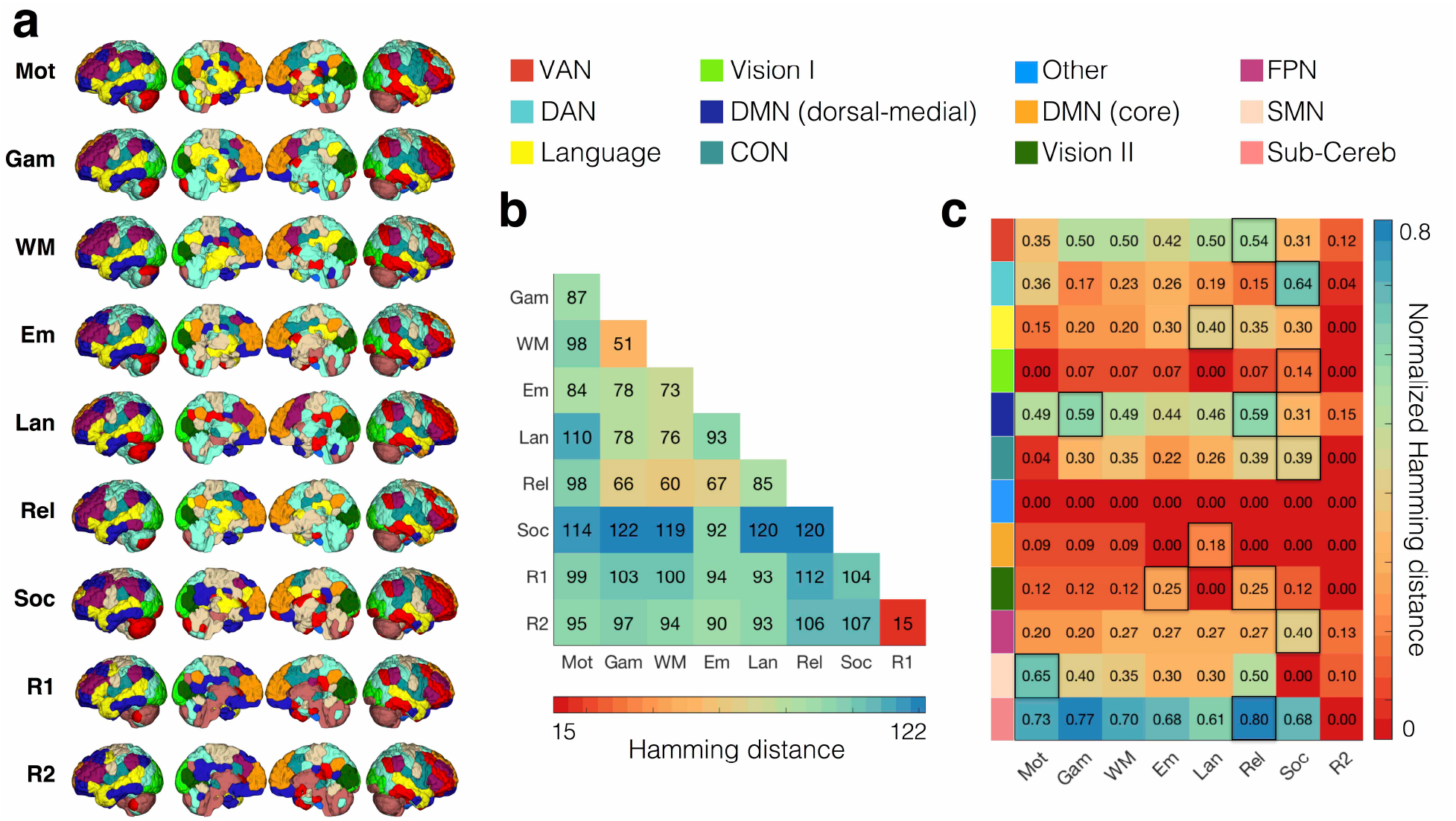
Functional network configuration varies across states. a) We computed the state-specific population-level node-to-network assignment (NNA) as the winner-takes-all assignment for each node across all individuals for each functional state (MOTOR, GAMBLING, WORKING MEMORY, EMOTION, LANGUAGE, RELATIONAL, SOCIAL, REST1, and REST2). b) Hamming distance between every pair of parcellation schemes derived for each state. The Hamming distance is calculated as the number of nodes that are assigned to different networks between the two functional states. It takes values from 0 (perfect matching) to the number of nodes (here, 268). As expected, the two resting states have the most similar parcellations (minimum Hamming distance: 15/268 = 5.6%). SOCIAL and GAMBLING had the least similar parcellations (minimum Hamming distance: 122/268 = 45/5%). SOCIAL task displayed the least similarity to the rest of the states with 41.9% (=112.25/268) different assignments on average. c) The ratio of the nodes that changed their network assignments from REST1 to every other state, calculated for every network separately. For every network, the state during which the maximum reconfiguration is observed is highlighted in black. VAN: ventral attention network, DAN: dorsal attention network, Vision I: primary vision, DMN: default mode network, CON: cingulo-opercular network, Vision II: secondary vision, FPN: frontoparietal network, SMN: sensorimotor network, Sub-Cereb: subcortical/cerebellum. See Figure S1 for a probabilistic illustration of these network definitions.

We next quantified the level of state-evoked reconfiguration for each network by computing the ratio of the nodes that change their network assignments from REST1 to every other state (Figure 1c). Among all eight states, REST2 exhibited the maximum similarity to REST1, with 7/12 networks displaying 0% reconfiguration. The subcortical/cerebellum exhibited the maximum reconfiguration, while the visual networks (visual I and II), and the DMN (core) exhibited the least reconfiguration from REST1 to all other states. The SMN displays the maximum reconfiguration during MOTOR task, and the language network displays the maximum reconfiguration during LANGUAGE task. Similarly, a majority of the higher-order association networks—including the FPN, DAN, and CON—display their maximum reconfiguration during the SOCIAL cognition task (Figure 1a). These observations could indicate that specific network reconfigures when a task strongly engages that network. These fuzzy network profiles are displayed in Figure S1 (see Methods).

### Decoding functional states by node-to-network assignment (NNA) vectors as features

Next, we demonstrate that the observed state-evoked reconfiguration of functional networks is robust across subjects and specific to that state. To this end, we trained a fully cross-validated predictive model using a gradient boosting machine (GBM) to predict the states of novel subjects based solely on their NNAs. We randomly divided the entire population into a training set and a test set. We calculated exemplars using the training set and used to assign nodes to networks for the entire population. Next, we trained a GBM on the NNAs of the training set and predicted the state for the test set subjects.

We report two classification pipelines: a two-class (binary) classification (Figure 2a), where every pair of states were used versus each other; and a more difficult 8-class classification (Figure 2b), where all states were used together with the exception of REST2 due to its overlapping nature with REST1. Random accuracy is 50% for the binary classification and 12.5% for the 8-class classification. As expected, the minimum prediction accuracy was associated with the REST1-REST2 pair (accuracy=55%). We successfully decoded all other pairs of states with accuracies considerably higher than random, with minimum, maximum, and average accuracy of 84%, 97%, and 94% averaged over all Ks from 2-50 (Figure 2a). Similarly, the 8-class classification accuracies were also considerably higher than random with minimum, maximum, and average accuracy of 66%, 87%, and 78% averaged over all Ks from 2-50 (Figure 2b). Together these predictive models suggest that functional networks reorganize due to changes in state in a robust, reliable, and predictive manner across subjects.

**Figure 2.**
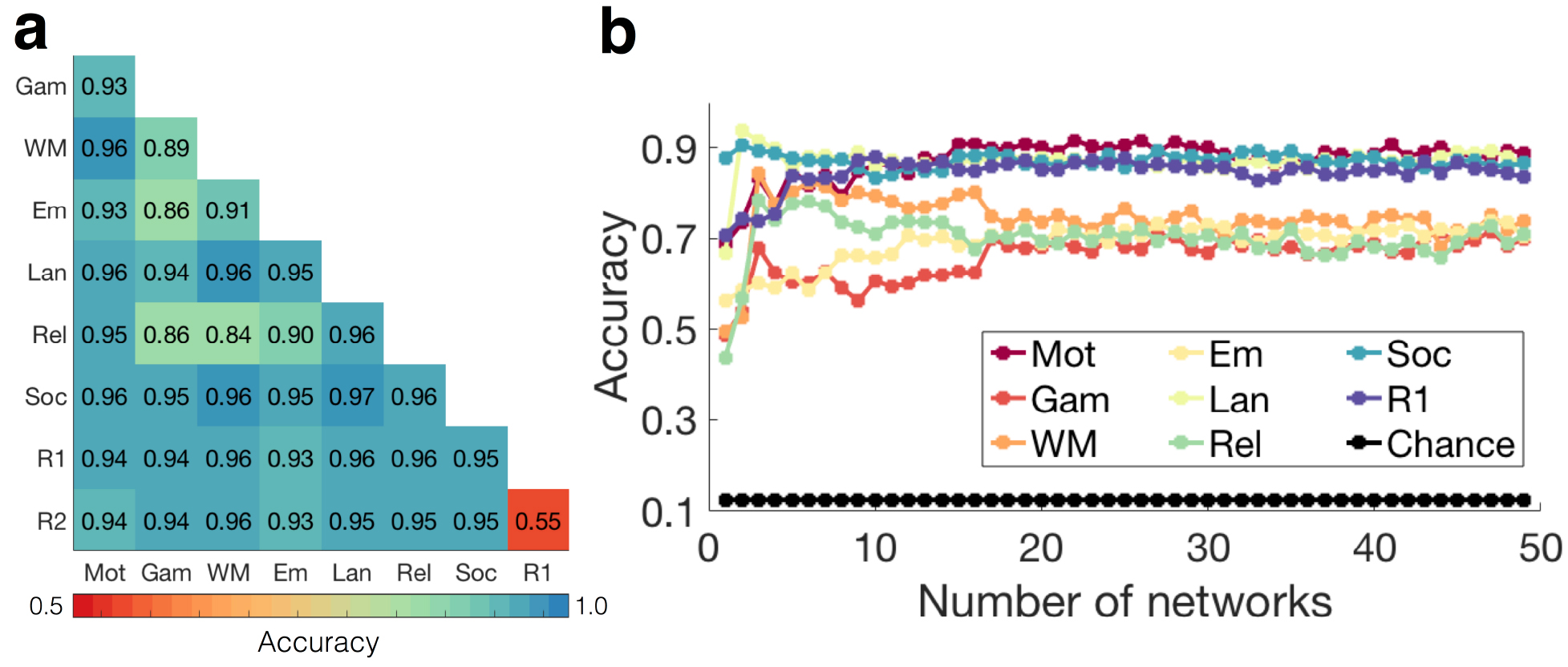
Decoding functional states by node-to-network assignment (NNA) vectors as features. The entire population was divided into equal-sized training and testing sets. The exemplars (nodes with fixed NNAs both across states and subjects, representing each network) were computed using the training set and were used to parcellate the entire population. A GBM was trained on the training set using the NNA vectors as features. It was then used to predict the functional state of the novel subjects in the testing set. a) The binary classification accuracies averaged over number of networks ranging from 2 to 50. For every pair of tasks (task1, task2) we restricted our analysis to the data derived from the two tasks, yielding a population of 718 × 2 subjects with two outputs, task1 or task2. b) The 8-class classification accuracies are displayed for all number of networks (K=2-50). For Ks larger than 17 the accuracies tend to stabilize with marginal variation. As expected, the minimum accuracies are observed for K=2, since the entire vector only consists of two numbers: 1 or 2, depending on whether the node is assigned to network #1 or network #2. GAMBLING, RELATIONAL, EMOTION, and WM displayed the lower prediction accuracies than MOTOR, SOCIAL, LANGUAGE, and REST. See Figure S4 for a 10-fold cross-validated predictive model wherein the parcellation schemes are derived from the entire population (not just the training set as shown above).

### Classifying nodes based on node-to-network assignment (NNA) reconfigurations

To better quantify which nodes change their NNAs across states and subjects, we calculated the entropy (a well-established metric of uncertainty from information theory literature (39)) of the NNA histograms. We used two measures of entropy: 1) Entropy_cross–subject_ to measure the variation across multiple groups of subjects within the same state (i.e. within a column) and 2) Entropy_cross–state_ to measure the variation across multiple states for the same subject (i.e. within a row). A high entropy in either of these measures means high variation in NNAs. Figure 3a displays Entropy_cross–state_ versus Entropy_cross–subject_ for every node in the brain. We observed that the two entropies were significantly correlated with each other (r=0.78, p<1×10^-16^), suggesting that the nodes with high cross-subject variation also tend to have high cross-state entropy.

**Figure 3.**
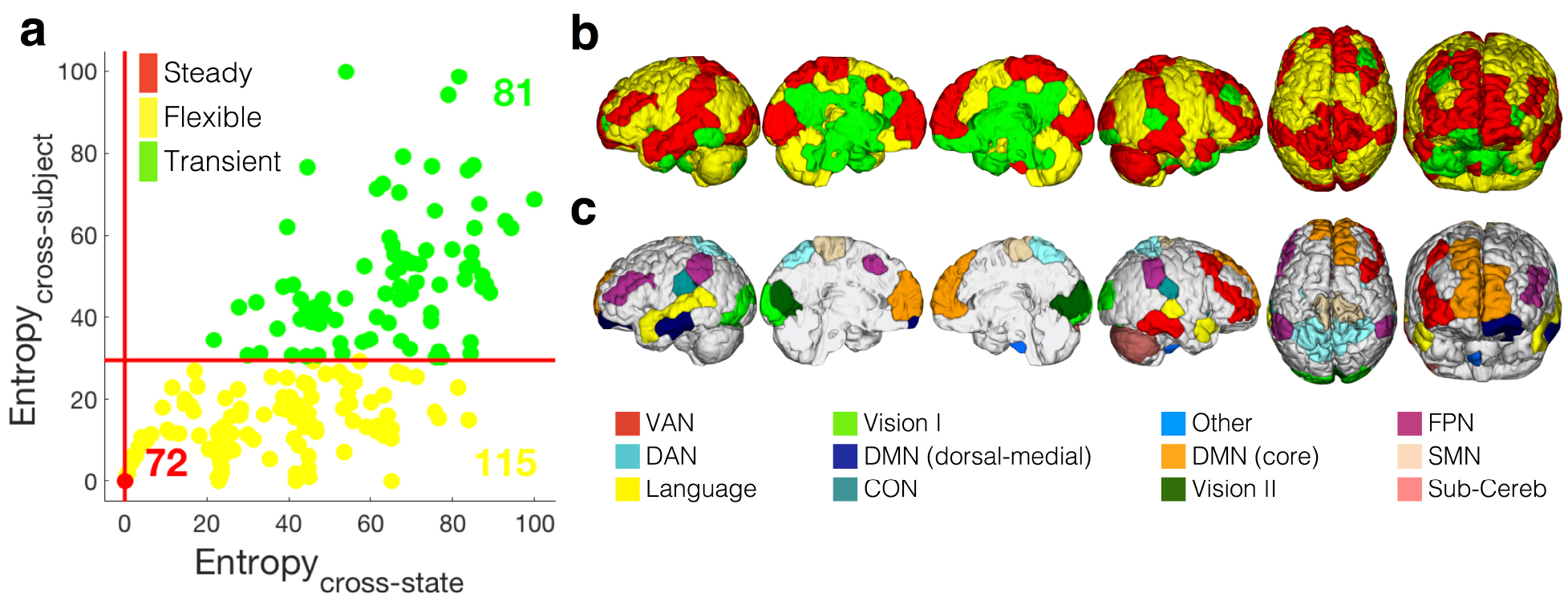
Classifying nodes based on node-to-network assignment (NNA) reconfigurations. a) For every node, we calculated two measures of variability: the node-to-network assignment (NNA) entropy across states (Entropy_cross-state_) and the NNA entropy across subjects (Entropy_cross-subject_). The two measures are significantly correlated (r=0.78, p<1×10^-10^). We grouped nodes into three entropy classes: stable nodes (red, n=72/268) have highly consistent network assignments both across states and across subjects; flexible nodes (yellow, n=115/268) change their network assignments across different states, but these changes in network assignments are consistent across subjects; transient nodes (green, n=81/268) change their network assignments both across states and sessions. b) The three entropy classes are visualized on the brain. The steady class includes areas in superior frontal cortex, a large portion of the temporal and occipital lobe. The flexible class revolves around higher order association areas including a large portion of the frontal and parietal lobes. The transient class is mainly localized in the subcortex and cingulate cortex. See also Figure S5 for a more detailed illustration of these entropy classes at the population and subject level.

Next, we used the two entropy measures to categorized nodes into three entropy classes: 1) *steady* nodes, those with low cross-state and cross-subject entropies, indicating consistent cross-state and cross-subject NNAs; 2) *flexible* nodes, those with relatively high cross-state entropies, but low cross-subject entropies, indicating inconsistent cross-state NNAs but consistent cross-subject NNAs; and 3) *transient* nodes, those with high cross-state and cross-subject entropies, indicating inconsistent cross-state and cross-subject NNAs (see Figure 3a).

The first class includes nodes that have consistent NNAs across subjects and across states. These nodes have low values for both entropy measures defined above, and are denoted as *steady* nodes (displayed in red in Figure 3a). The second class includes nodes that have consistent NNAs across subjects, but that change their NNAs across states. These nodes have lower values for Entropy_cross–subject_, but higher values for Entropy_cross–state_ and are denoted as *flexible* nodes (displayed as yellow in Figure 3a). The third class includes nodes that change their NNAs across different states and subjects. These nodes have high values for Entropy_cross–subject_ and Entropy_cross–state_ and are denoted as *transient* nodes (displayed as green in Figure 3a). As shown in Figure 3b, steady nodes are mainly located in the superior frontal cortex, dorsolateral prefrontal cortex (dlPFC), precuneus, post-central gyrus, superior and middle temporal gyrus, and the occipital lobe. Flexible regions are located mainly in the inferior frontal gyrus, pre-central gyrus, anterior cingulate cortex, posterior parietal cortex, inferior parietal lobule, and cerebellum. Finally, transient nodes are located mainly in subcortical areas such as the hippocampus/parahippocampus, thalamus, and caudate. Figure 3c illustrates network assignments for the steady nodes. These results suggest that the brain is comprised of nodes that form three general entropy classes based on NNA: steady, flexible, and transient.

### Localization of the node entropy classes in networks

We investigated the association between the three node entropy classes and our functional networks. We observed that all networks—excluding the visual I network—contained nodes of each class. However, the distribution of the three entropy classes differed considerably across networks (Figure 4a). For example, the DAN (76.6%), CON (56.5%), and SMN (70.0%) had a majority of flexible nodes; the VAN (57.7%), and the dorsal-medial portion of DMN (64.1%) had a majority of transient nodes; and the visual I network (62.5%), visual II network (64.3%), and the core portion of DMN (72.7%) had a majority of steady nodes. Finally, we observed that all functional networks contributed evenly to the steady class, but did not contribute evenly to the flexible or transient classes (Figure 4b). For example, the DAN contributed the most to the flexible class, while the dorsal-medial portion of DMN and the subcortical/cerebellum contributed the most to the transient class. This suggests that individual brain networks have varying levels of stability, as defined by NNA entropy across task states and subjects.

**Figure 4.**
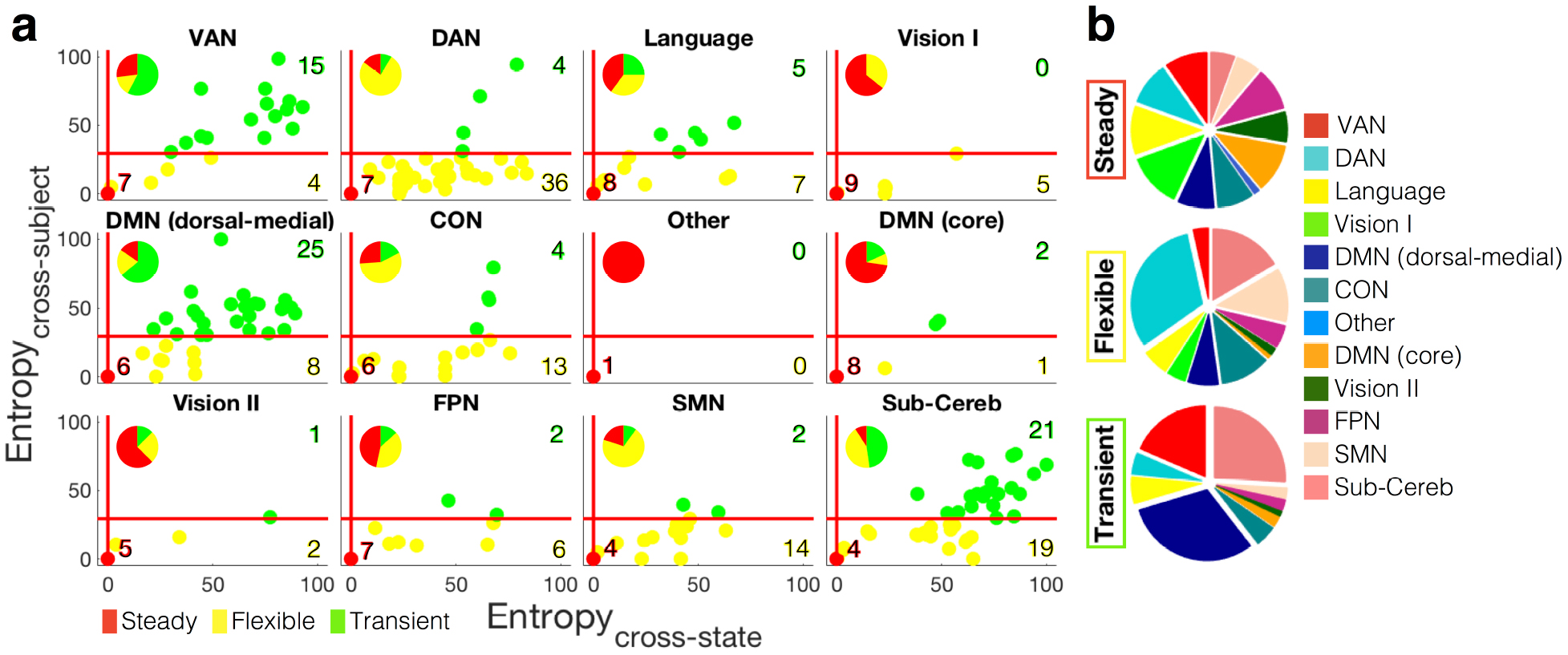
Localization of the node entropy classes in networks. a) The Entropy_cross-state_ versus Entropy_cross-subject_ diagram is displayed for the nodes within each network. The within-network distribution of the entropy classes is further demonstrated. Primary and secondary visual networks and the core portion of default network are mainly comprised of steady nodes. DAN, CON, and SMN mainly include flexible regions, and VAN, dorsal-medial DMN and subcortical/cerebellum networks are mainly comprised of transient regions. FPN has a large portion of both steady and flexible regions, subcortical/cerebellum network has a large portion of both flexible and transient nodes, and Language network has almost even portion of all three entropy classes. b) The distribution of functional networks within each entropy class. While the contribution of functional networks to the steady class is evenly distributed across all networks, DAN, CON, SMN, and subcortical/cerebellum have the largest contribution to the flexible class, and VAN, dorsal-medial DMN, and subcortical/cerebellum have the largest contribution to the transient class.

### Flexible nodes contributed most to the state-decoding paradigm

We hypothesize that nodes in the flexible entropy class should have the highest predictive power for decoding states. We support this hypothesis with two lines of logic: first, because flexible nodes change their network organization according to task state, how the NNA reconfigures should therefore reflect the underlying state; second, because (as defined above) flexible nodes have consistent NNAs across subjects, NNA should generalize across novel subjects. To test our hypothesis, we attempted to predict task state from NNA and observed how important each NNA was in this prediction. Node importance (i.e. “feature importance”) was calculated by the GBM classifier as the number of times that each node was used to make a key decision that improved the classifier’s performance measure (40). We examined the distribution of the importance scores within each node entropy class. Consistent with our hypothesis, the flexible regions had the largest importance scores on average (Figure 5a; two-tailed t-test Bonferroni corrected for multiple comparisons, t(flexible, steady)=26.8, p<3×10^-16^, t(flexible, transient)=30.4, p<3×10^-17^, t(steady, transient)=0.34, p<0.79). This suggests that entropy classes are functionally meaningful divisions and that task-state can be decoded based on the NNA of the brain’s flexible nodes.

**Figure 5.**
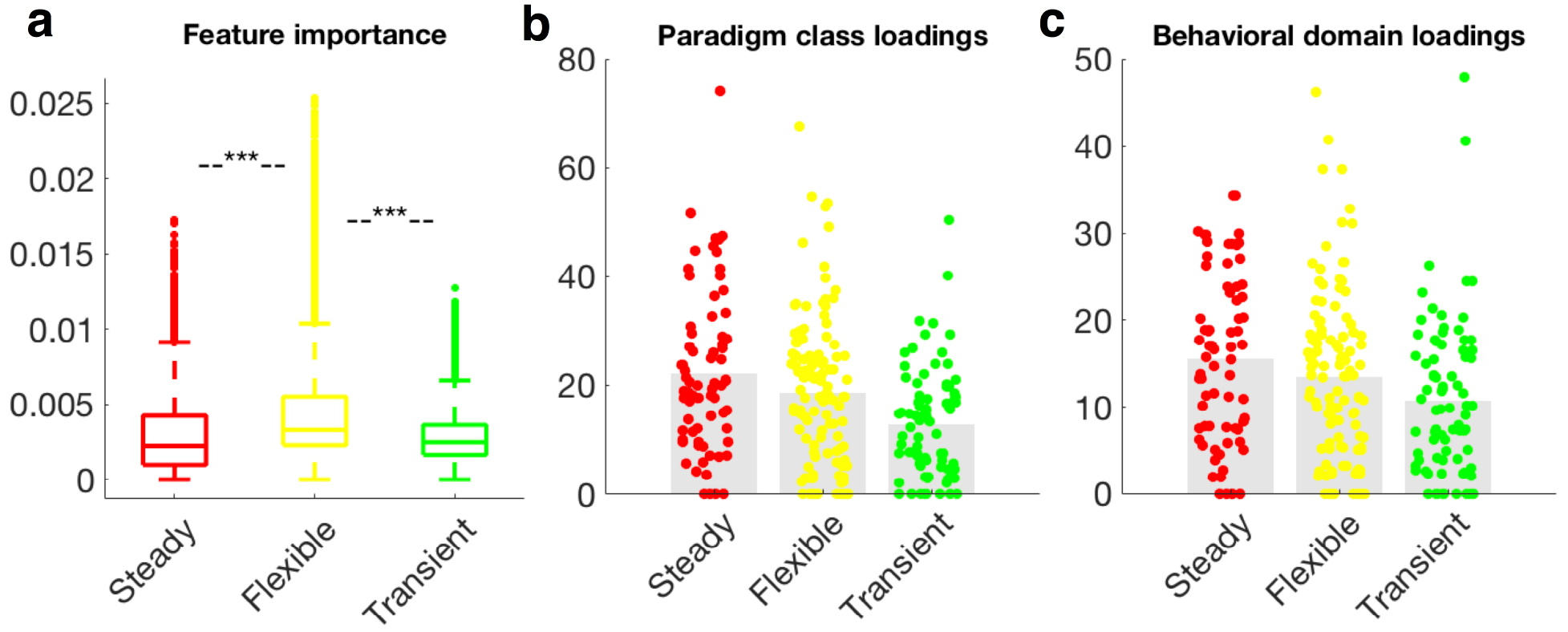
Predictive power and functional relevance of the node entropy classes. a) The distribution of the feature (node) importance in predicting states is displayed across the three entropy classes. The flexible class comprises of the most important nodes for state decoding. Two-tailed t-test between every pair of steady, flexible, and transient class was performed and Bonferroni corrected for multiple comparisons, *** p<1×10^-16^. b) The distribution of paradigm classes (PCs) across entropy classes. Steady nodes have higher values of significant PC loadings than either flexible (t=1.7, p<0.09) or transient nodes (t=4.6, p<8×10^-6^). c) The distribution of behavioral domains (BDs) across entropy classes. Steady nodes have higher values of significant BD than either flexible (t=1.5, p<0.15) or transient nodes (t=3.3, p<0.001). Analyses in (b) and (c) represent node-level analyses of BrainMap Functional Database’s experimental meta-data representing 55 BDs and 108 PCs. Profiles were computed within each node as the forward inference likelihood z-scores, thresholded at z>1.96, and summed over all BD/PCs. Colored circles represent the 268 nodes, colored according to their entropy classes. See Figure S6 for network-level functional relevance.

### Steady nodes displayed the strongest behavioral associations

Similarly, we investigated the behavioral profiles of different entropy classes. We conducted a meta-analysis using the behavioral domains (BD) and paradigm classes (PC) reported in the BrainMap Functional Database (41). We examined the association between BrainMap’s 55 BDs and 108 PCs (reflecting 10,467 task-activation experiments from 27,820 subjects) with the node entropy classes (steady, flexible, and transient), both at the node-level (with 268 nodes) and at the network-level (with 12 networks). We hypothesized that because steady nodes were less likely to “switch” association with task state, they should be more strongly associated with specific BDs or PCs. We found that indeed, there was a trend for steady nodes to have stronger BD and PC loadings than either flexible or transient nodes (One-tailed t-test Bonferroni corrected for multiple comparisons, PC: t(steady, flexible)=1.7, p<0.09, t(steady, transient)=4.6, p<8×10^-6^; BD: t(steady, flexible)=1.5, p<0.15, t(steady, transient)=3.3, p<0.001; Figure 5b,c). The same trend was observed at the network-level (Figure S6). A breakdown of each node’s BD and PC association is referenced in Table S1. This further suggests that entropy classes provide information that is relevant to how the brain executes its functional repertoire, as defined by BDs and PCs.

### Graph visualization of the functional connectivity reconfiguration across states

Finally, we calculated changes in functional connectivity due to changes in state, for comparison with studies that investigate changes in functional connectivity as a proxy for network reorganization. Functional connectivity matrices were calculated (using Pearson correlation), z-scored, and averaged across subjects. The averaged connections were then thresholded to the top 10% of the connections. Figure 6 (the right side of each panel) visualizes the resulting connectivity matrices as force-directed graphs with nodes colored according to the functional network to which they belong and edges colored according to the entropy class of their two end points (see Methods). These force-directed graphs aim to visually organize networks such that the energy of the graph as a whole is minimized. This is accomplished by assigning both repulsive and attractive forces to each pair of nodes such that the nodes with stronger interconnections are displayed closer to each other and the ones with weaker connections are depicted distant. Node size is proportional to the graph theory measure degree. Nodes from different but related networks were integrated for all states. For instance, the core and dorsal-medial portions of DMN as well as the visual I and visual II networks showed strong integration with each other. However, the magnitude of this integration differed across different states. For example, while the visual I and visual II networks displayed strong integration across all states, the functional connectivity between them was weaker during the Social task. Finally, networks changed their segregation and integration patterns as a function of state. For instance, the DAN differed considerably across states, with higher segregation during the task states and higher integration during the rest states. Conversely, the subcortical/cerebellum and SMN presented higher segregation during the rest state and higher integration during the task states.

**Figure 6.**
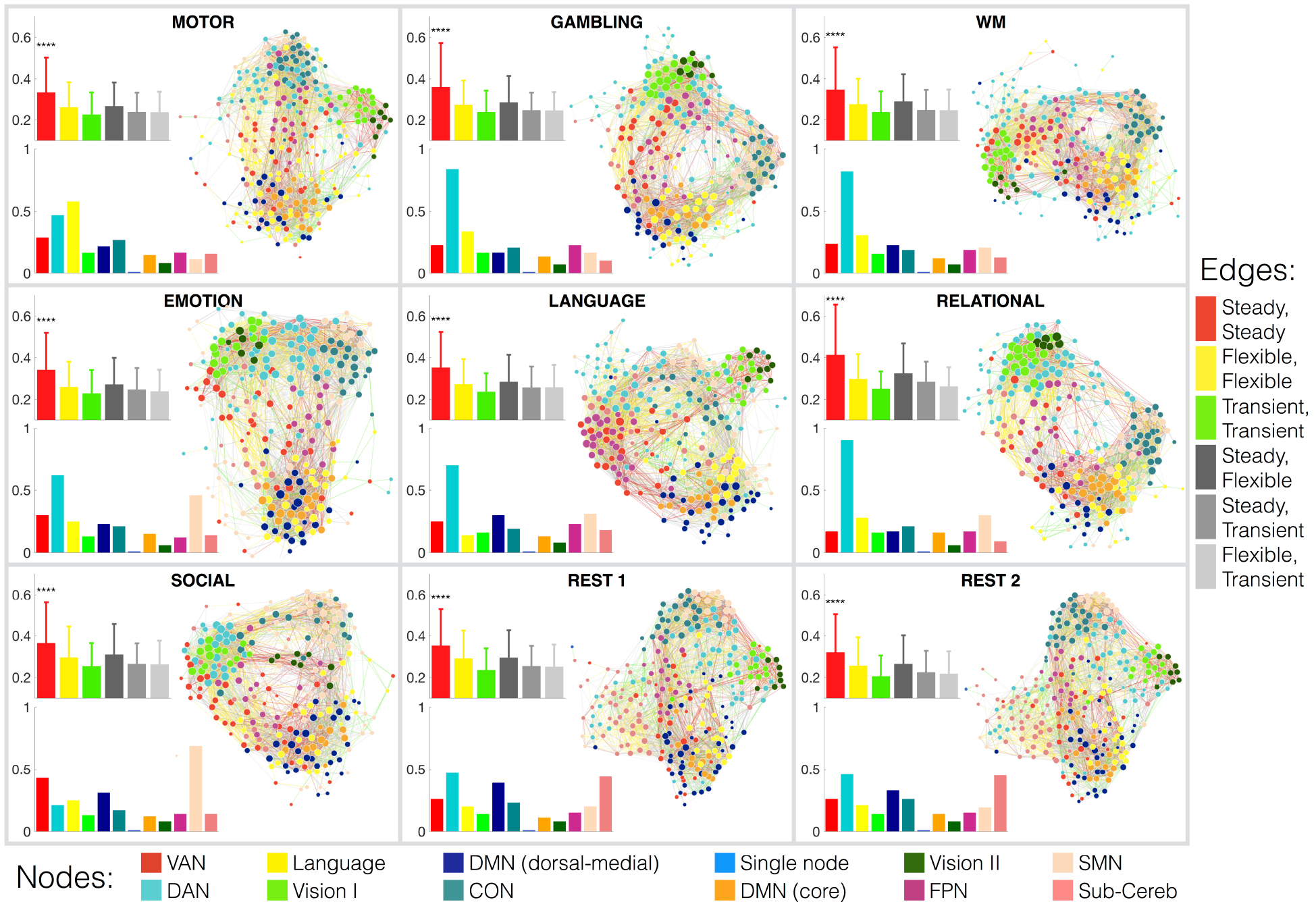
Graph visualization of the functional connectivity matrices across functional states. Each panel corresponds to one functional state. For each functional state, three diagrams are displayed. The right diagram is the graph visualization of the functional connectivity matrix averaged over all subjects. Nodes are colored according to their population-level network assignments (see the bottom legend bar), and are sized based on their degree values (the higher the degree of the nodes, the larger it is displayed). Edges are colored according to the entropy class of their two end points (see the right legend bar), for example the connections between two steady nodes are colored red, connections between a steady node and a flexible node are colored light gray, and so on. The graphs are structured such that nodes with stronger connections are spatially closer to each other. As expected, nodes within the same network are clustered spatially close to each other. Further, nodes from different but related networks are also spatially integrated (for example the core and dorsal-medial portions of DMN, or primary and secondary visual networks). However, there is a difference in the integration and segregation level of different functional networks across different task states. The bottom left diagram within each panel represents the ratio of the nodes assigned to each functional network defined at the population-level. The top left diagram within each panel represents the strength of different types of connections at the population-level. Error bars represents the standard deviation across connections. Surprisingly, the connections within steady nodes are significantly stronger than the rest of the connections. These steady nodes form a strong core organization in the brain with large interconnections, similar to the “rich club” pattern.

We next quantified the reorganization of functional networks by exploring their distribution across states (Figure 6; the bar plots on the bottom-left of each panel). We observed that the distribution of functional networks changes considerably across states, with more uniform distribution during rest than task. In this sense, the DAN was the largest network (with the maximum number of nodes) during GAMBLING, WM, EMOTION, LANGUAGE, and RELATIONAL states. The subcortical/cerebellum network was considerably larger during rest than task, suggesting that nodes in the subcortical/cerebellum network tend to integrate with other networks during the execution of a task, while displaying segregated behavior during rest (consistent with our observation from the graph). Surprisingly, the SMN lost the maximum number of nodes during the MOTOR task, while the Language network lost the maximum number of nodes during the LANGUAGE task. While this observation may seem contradictory to the demands of the tasks, it in fact indicates that these networks tend to integrate most with other networks during these tasks, potentially facilitating the information flow across networks, an observation aligned with the literature (22, 42). This is also consistent with our earlier finding (Figure 1c) that these networks displayed the maximum reconfiguration during these tasks.

Next, we quantified the functional connectivity (i.e. edge strength) between nodes from the same or different classes (Figure 6; the error bars on the top-left of each panel): steady-steady edges (colored as red), steady-flexible edges (colored as dark gray), steady-transient edges (colored as gray), flexible-flexible edges (colored as yellow), flexible-transient edges (colored as light gray), and transient-transient edges (colored as green). We observed that the edge strength between steady nodes was significantly stronger than the edge strength between other node classes (p<2×10^-16^, Bonferroni corrected for multiple comparisons). Among edges between nodes of the same class, edges between steady nodes were significantly stronger than the edges between flexible nodes (p<2×10^-16^, Bonferroni corrected for multiple comparisons) and edges between flexible nodes were significantly stronger than edges between transient nodes (p<1×10^-2^, Bonferroni corrected for multiple comparisons). Among edges between nodes of differing class, the edges between steady and flexible nodes were the strongest (p<2×10^-2^, Bonferroni corrected for multiple comparisons). Edges associated with transient nodes had the lowest strength.

## Discussion

In this work, we showed that the brain’s network organization is not fixed, but rather that networks reconfigure as a function of brain state. We showed this held true across different task states and across individuals. We measured network reconfiguration as node-to-network assignments (NNA) and demonstrated that a novel subject’s current state could be predicted with 66-97% accuracy based on NNA alone. This finding demonstrates that networks reconfigure in a meaningful and reproducible manner across brain states. This also highlights the robustness of state-evoked network reconfigurations across subjects. Further, we showed that nodes group into three entropy classes based on their NNAs across states and subjects. Steady nodes exhibit consistent NNAs across both states and subjects and were primarily located in visual and medial cortical regions. Flexible nodes change their network assignments according to state in a consistent manner across subjects and were primarily located in higher order cognitive regions. Transient nodes exhibit variable NNAs across both states and subjects and were primarily located in the sub-cortex and cerebellum; regions known to have lower reliability in functional connectivity studies (43). Despite these trends of anatomical locations of nodal classes, all networks contained some nodes from each class. We further demonstrated the functional relevance of these nodal classes: flexible nodes contribute the most to models predicting brain state and steady nodes display the largest behavioral loadings based on a large-scale meta-analysis of task activation studies. Together, these results demonstrate that the brain’s functional networks recruit or dismiss individual nodes according to the demands of a particular task state.

### Networks reorganize during different task-states

Our results suggest that a similar core network structure is observed across many different brain states. The same classic resting-state networks were observed in all brain states, but the exact nodes that belong to these networks changed as a function of the specific state. These results suggest that the organization of nodes into networks is state dependent, that network structure is not fixed across task states. Furthermore, our BrainMap meta-analysis showed that nearly all nodes were significantly associated with multiple paradigm classes, arguing against a single NNA or a discreet functional mapping for a specific node. Altogether, our results suggest that NNA is inherently probabilistic (see Figure S1) as the brain is constantly reconfiguring network membership. As such, previous studies may have interpreted these changes in NNA as changes in functional connectivity between networks because they considered network membership as a fixed quality (44). Future studies investigating changes in functional connectivity between rest and task or between multiple tasks should account for these changes in network structure to fully characterize network dynamics.

Our key finding is that all networks reconfigure as a function of task. This contrasts with previous literature that has focused on a single network, e.g. the FPN, that changes as a function of task-state (25, 45). Another contrast, for example, is the SMN, which has been characterized as remaining relatively fixed (11). However, we observed that the SMN is highly variable across states (Figure 4a), revealing a state-evoked reconfiguration not previously characterized. Perhaps as one would expect, the largest reconfiguration of the SMN occurs when subjects are executing the motor task (Figure 1c). The SMN had the lowest portion of transient regions, suggesting the low cross-subject variability, consistent with previous observations (37, 46). In addition, our results also replicate previously observed network dynamics. Consistent with Andrews-Hanna et al. (47), we highlight that there is a core sub-system of the DMN (including medial superior frontal gyrus) and dorsal-medial and medial-temporal sub-systems of the DMN (including ventral medial prefrontal cortex (vmPFC), inferior temporal gyrus and temporal pole). Our results extend this characterization by suggesting that the two sub-systems of the DMN display distinct dynamic behaviors, with the core subsystem remaining unchanged across states, and the dorsal-medial subsystem reconfiguring more flexibly across states and subjects (48). Highlighting the complex role of DMN in both tasks and rest, our observation indicates that the DMN nodes change their network assignments from the DMN to other networks (such as the FPN and CON). Overall, these observations support the notion that all networks reorganize due to task, likely in a task specific manner.

### Networks contain steady, flexible, and transient nodes

Our findings reveal that all networks contain a combination of steady, flexible, and transient nodes (Figure 4a). This suggests that networks consist of a core set of steady nodes that then recruit and dismiss flexible and transient nodes as required by a particular task demands. This result further implies that within each network there are specialized regions associated with distinct functionalities, from maintaining the intra-network communication to expanding the inter-network integration. These functionalities are corroborated by the theory of “local” versus “distributed” neural communication (22, 42), where transitions across states require the segregated processing units (distinct functional networks) to integrate information through flexibly changing network membership.

The contribution to the steady nodes was evenly distributed across all networks (Figure 4b), indicating all networks retained some core functional network configuration. The majority of steady nodes were located in regions previously been implicated as part of the “rich club”, a common organization in complex systems where important (or “rich”) nodes connect preferentially to other important nodes (15, 49, 50). Additionally, we observed strong functional connectivity between nodes in the steady class (Figure 6, the top left error bars) in a similar vein to the connections between rich club nodes. These observations were consistent across all nine states, implying that the “rich club” organization of the brain is independent of task-state. This also mirrors a core-periphery structure in the brain with core nodes strongly and mutually interconnected and periphery nodes sparsely connected to each other and to the core nodes (50-52).

In contrast, the distribution of flexible nodes varies across networks, suggesting that some networks reconfigure more with brain state changes than others (Figure 4b). For example, the DAN, CON, and FPN had a larger proportion of flexible nodes (Figure 4a,b). Previous studies report that these networks rapidly update their connectivity patterns according to the task context (19, 20, 25, 53, 54) facilitating the information flow across networks (22, 42). The large number of nodes that change their network assignments to the DAN, CON, and FPN may underlie these previously observed changes in connectivity patterns. Our results are also consistent with the role of these networks as functional hubs (25, 55) and their strong out-of-network connections (42).

Finally, the majority of the transient nodes were assigned to the subcortical/cerebellum, dorsal-medial subsystem of DMN, and VAN (Figure 4a,b). These nodes, mainly located in heteromodal association and limbic cortex, actively change their membership across both subjects and states. High cross-subject variability in these regions has been previously reported (46, 48), and linked to differences in intelligence (56), attention (57, 58), and personality traits (59). This suggests that these regions promote a more personalized reconfiguration in the brain to adapt to the task at hand. Future work could explore the extent to which differences in the network reconfiguration in these regions could be indicative of, or be inferred from, the individual differences in task performance.

### Network configuration predicts state

Individual task-states can be predicted from NNAs, suggesting that the state-specific information of network organization is sufficiently robust and reliable to form a signature for a given state. Prior work on task decoding has typically employed binarized classification of tasks. The pair-wise accuracies (binarized classification) achieved here (ranging from 84% to 97% with average of 94%; Figure 2a) are significantly higher than the ones reported in the literature (60-62). Our accuracies for the 8-class classification (ranging from 66% to 87% with average of 78%; Figure 2b) were considerably higher than the accuracies reported in the literature, even for easier decoding problems such as classification with lower number of classes (25) or within-subject classification paradigms (63). In contrast to previous approaches that use voxel-wise feature (63, 64)—and either suffer from the “curse of dimensionality” or heavily rely on *a priori* set of voxels—our prediction results use a whole-brain data-driven feature engineering approach. That the minimal information stored in the NNA vectors (of length 268) with only integer values (i.e. 1, …, K for K networks) can predict the state for novel subjects highlights the amount of state-specific information stored the brain’s large-scale network organization.

### Individualized brain networks are needed

In addition, our results highlight the need for individualized approaches to systems neuroscience and medicine (65, 66) by demonstrating that NNAs contain large individual variations across states (i.e. our transient node class). Individualized, state-specific networks may provide higher specificity when considered as predictive features in machine learning algorithms (67, 68). Individualized networks could provide input for real-time fMRI neurofeedback paradigms (69-71), and brain stimulation therapies (72, 73), focusing on the disruption of networks at the single subject level. The conventional approach of defining fixed networks via group-level parcellations could reduce the efficacy of treatments.

In a recent work, Gratton et al. have studied the magnitude of functional network variability across subjects, tasks, and sessions (29). Their study of brain reconfiguration was restricted to changes in functional connectivity matrices as a whole (i.e., correlation of the vectorized connectivity matrices). Our work differs in that we study the large-scale network reconfigurations by probing the changes in NNAs across subjects and states. These two divergent but complementary views shed light on different aspects of the brain dynamics. While their analysis has shown that functional connections are largely stable with subtle cross-state changes (in terms of correlating vectorized matrices), here we demonstrate that about 73% (196/268) of the nodes actually change their network membership as a function of state. In addition, their observations of functional connectivity matrices indicate that the state-evoked modulations are largely individual-specific (i.e., subtle state-specific changes are consistent across subjects), whereas we demonstrate that state-specific changes in NNAs are sufficiently robust and reliable across subjects to predict the brain state. Combined, these observations suggest that while gross matrix level connectivity patterns change only minimally with state, large-scale NNAs change in a significant and meaningful manner. Therefore, NNA is state-sensitive and can summarize whole-brain connectivity across different task states. Our results show that the network level organization of the brain is flexible and changes in meaningful ways as a function of brain state, even at the individual level.

### Further considerations

This work has several limitations that should be noted. Although brain-state dependent network reconfigurations were investigated here, we did not investigate the temporal dynamics within a given condition. This was partly because the length of the sessions in HCP data vary (74, 75) and because of methodological concerns surrounding dynamic brain-state definitions. Previous studies using window-based approaches to capture brain dynamics have shown sensitivity to the length of the window employed (23). We suggest that future work extend our framework to study the within session dynamics by employing sliding windows of sufficient length.

We employed a functional atlas consisting of 268 nodes, generated from an independent group of healthy subjects (5, 6). This atlas has proven to be reproducible and reliable (5), beneficial in understanding cross-subject variability, and useful for developing predictive models of behavior (6, 76-78). Given this past performance, this 268-node atlas represents a reasonable way to operationalize the brain’s sub-units in network analyses; however, we expect these results to generalize to any generic node-level atlas of comparable resolution. It should be noted however, that while we have assumed a fixed node definition across subjects and states (as have most network level studies to date), the same notion of flexibility is expected at the node level, and we are currently extending this approach to examine node boundary reorganization as a function of task or brain state.

Finally, moving forward it will be important to relate state-evoked network reorganization with individual differences in behavior and/or clinical measures. Future work could quantify the extent to which subjects with similar NNA reconfiguration patterns are also similar in state, behavioral, or clinical measures; potentially grouping homogeneous subjects based on their NNA phenotypes. This methodology could be particularly compelling when applied to subjects with different clinical, neurodevelopmental, and aging categories.

### Conclusion

We showed that the brain’s network organization is not fixed, but rather that networks reconfigure in predictable ways as a dynamic function of brain state. Future work should consider these state-evoked reconfigurations when studying changes in connectivity during different task-states. Such an approach will hopefully allow us to begin to relate individual differences in network reconfiguration to individual differences in behavior or clinical symptoms.

### Code availability

The 268-node functional parcellation is available online on the BioImage Suite NITRC page (https://www.nitrc.org/frs/?group_id=51). MATLAB, R (for graph visualization), and Python (for predictive modeling) scripts were written to perform the analyses described; these code are available on GitHub at https://git.yale.edu/ms2768/NetworkParcellation_States.git. The graph visualization is released separately under the terms of GNU General Public License and can be found here: https://github.com/mehravehs/NetworkVisualization.

## Acknowledgments

Data were provided by the Human Connectome Project, WU-Minn Consortium (Principal Investigators: David Van Essen and Kamil Uǧurbil; 1U54MH091657) funded by the 16 NIH Institutes and Centers that support the NIH Blueprint for Neuroscience Research; and by the McDonnell Center for Systems Neuroscience at Washington University. This work was supported by NIH Grant MH111424 and DARPA Young Faculty Award (D16AP00046).

## Author Contributions

M.S., A.K., D.S., and R.T.C. conceived and formulated the study. M.S. and A.K. developed the exemplar-based submodular parcellation algorithm. M.S. performed the parcellation analysis, graph visualization, and predictive modeling. D.B. and M.S. performed the behavioral mappings. M.S. wrote the manuscript with contributions from D.B., D.S., A.K., and R.T.C.

## Materials and Methods

### Participants and processing

Data were obtained from the 900 Subjects release (S900) data set in Human Connectome Project [HCP] (74). Analyses were limited to 718 subjects for which data were available for all nine functional states: MOTOR, GAMBLING, WORKING MEMORY (WM), EMOTIONAL, LANGUAGE, RELATIONAL, SOCIAL, REST1, and REST2. For details of scan parameters, see Uǧurbil et al. (75) and Smith et al. (79). Starting with the minimally preprocessed HCP data (80), further preprocessing steps were performed using BioImage Suite (81) and included regressing 12 motion parameters (Movement_Regressors_dt.txt), regressing the mean time courses of the white matter and cerebro-spinal fluid as well as the global signal, removing the linear trend, and low pass filtering (as previously described in Finn et al. (6)).

### Functional distance and functional connectivity matrices

Functional matrices were assessed using a functional brain atlas (5) consisting of 268 nodes covering the whole brain; this atlas was defined using resting-state data from a separate population of healthy subjects (6). For every subject in each functional state, time courses for the two functional runs with opposing phase-encoding directions (left-right, “LR”, and right-left, “RL”) were concatenated and further used to generate a ground set *V* consisting of *N* = 268 vectors in *T*-dimensional space, where *T* indicates the length of scan session. To construct 268 × 268 symmetrical connectivity matrices, the Pearson correlation coefficients between the time courses of each possible pair of nodes were calculated and normalized using Fisher’s z-transformation. Each element of the functional connectivity matrix represents a functional connection, or edge, between two nodes. Functional connectivity matrices were calculated for each subject and each state separately. We also constructed functional distance matrices to perform the network-level parcellation. To this end, all the data points were first normalized to a unit norm sphere centered at origin, and a point with the norm greater than two (*𝒱*_0_ = [3, 0, …, 0]) was used as the auxiliary exemplar. The pairwise squared Euclidean distances were calculated between the data points, yielding a distance matrix of size 268 × 268 for every subject and state.

### Individualized and state-specific functional network parcellation

Functional networks were delineated for every subject in each functional state using the exemplar-based parcellation algorithm developed earlier in our group (36). This method provides an approach to summarize data by introducing a set of *K* exemplars that best represents the full data. We followed the same methodology explained in Salehi et al. (36), by attempting to find *K* exemplar labels *S* = {*e*_1_, *e*_2_, …, *e_K_*} s.t. *e_i_* = ∈ {1, …, *N* = 268}, whose corresponding exemplar vectors for each individual *j* in each state *m*, i.e., 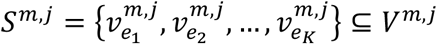, maximizes a desired utility function, as follows:

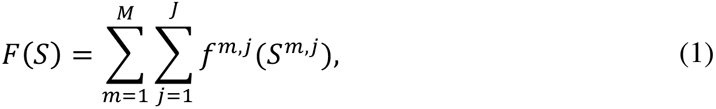

where *S* is the exemplar label set and *S^m^*^,*j*^ is the set including the corresponding exemplar vectors in subject *j* ∈ {1, …, *J* = 718} at state *m* ∈ {1, …, *M* = 9}. Here, 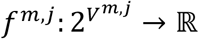 is the utility function of subject *j* at state *m*. This function is a nonnegative submodular function derived from a loss function *L* by introducing an auxiliary exemplar, *𝒱*_0_, according to equations (2-3):

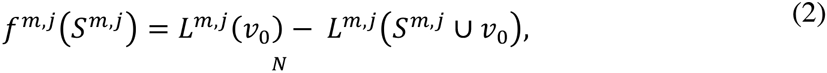

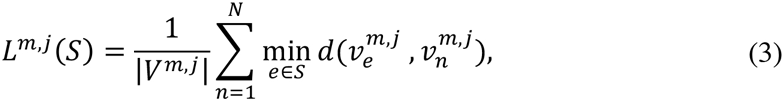

where *V^m^*^,*j*^ is the ground set of size 268 and dimension *T* for subject *j* at state *m*, and 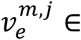 *S^m^*^,*j*^ ⊆ *V^m^*^,*j*^ is the node exemplar vector corresponding to the exemplar label *e* ∈ *S*. The greedy algorithm was employed to solve the optimization problem and find the best *K* = {2, 3, …, 50} exemplar labels.

After *K* exemplar labels were identified, the corresponding exemplar vectors in every subject and state were used to parcellate the brain. We assigned each region *n* in subject *j* and state *m* (i.e. vector 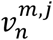) to the closest exemplar that represents the corresponding community (or network), i.e.:

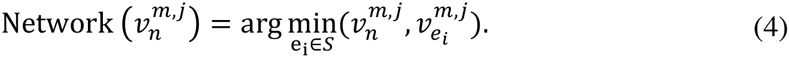

Thus, the brain was parcellated into *K* networks each represented by an exemplar.

Next, we obtained a group-level parcellation for each of the nine functional states. We employed the winner takes all algorithm over all subjects in each state. That is, region *n* was assigned to network *k* at state *m* if the majority of subjects in that state voted for this assignment.

### Local versus global exemplar set

Here, exemplars were identified such that they were common across all functional states (i.e., global exemplars), so that they are agnostic to the states that subjects occupy. This was to assure the proceeding predictive analyses were not biased by the prior knowledge of the state. In order to assure that the results were not confounded by introducing common exemplars across all states, we repeated the exemplar-based parcellation, this time restricting the analysis to each state separately. This approach yielded nine different sets of exemplars (i.e., local exemplars), one for each functional state. We compared the parcellation results from global and local exemplars using two separate measures. First, we calculated the Dice coefficient between the two group-level parcellation vectors (Figure S2a) for *K* = 2– 50. Second, we computed the Hamming distance between the community co-membership matrices derived from each parcellation. For every subject in each state, a network co-membership matrix (*A*) was constructed from the two parcellation schemes: (i) using global exemplars (*A*_global_), and (ii) using local exemplars (*A*_local_). The elements of this matrix were calculated as follows:

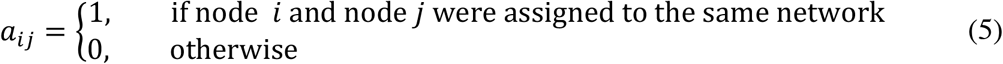

The similarity between the two co-membership matrices (*A*_global_ and *A*_local_) were calculated as 1 minus the Hamming distance between the vectorized form of the two matrices. The result is displayed in Figure S2b for *K* = 2 – 50.

### Number of networks (*K*) selected for the analysis

To choose the optimum number of networks, we examined the changes in network assignments by adding a new exemplar (or equivalently, adding a new network). For every state, we calculated the number of nodes that change their network membership as the number of exemplars, *K*, increases from 2 to 50. This was employed for every subject and state separately (Figure S3). Among the *K*s for which the number of changes were locally maximum, i.e. *K* = {12, 17, 31, 45}, we chose *K* = 12 as the minimum number of networks after which the memberships stabilized with significantly less changes in the NNAs. We reported the result for *K* = 12 in the main script and presented the analyses of the remaining *K*s in the Supplementary materials.

### The extent of network reconfiguration across states

We quantified the functional network reconfigurations across states by computing the pair-wise Hamming distances (38) between every pair of group-level parcellation vectors. This resulted in a distance matrix *D*_9×9_, where every element *d_m_*_1,*m*2_ represents the number of nodes that change their membership from state *m*_1_ to state *m*_2_. To study each individual network reconfigurations, we considered REST1 as the benchmark for comparison. For every network *k*, we computed the normalized Hamming distance between the NNA vector limited to the nodes that are assigned to network *k* during REST1, and the vector containing the same nodes’ network assignment during every other state.

### Functional state decoding

We established a fully cross-validated predictive model that predicts (or decodes) the functional state of each individual brain solely based on the NNAs. The predictive analysis was employed using two separate pipeliens: in the first pipeline, we employed a two-class (binary) classification task on every pair of states. For every pair of state *m*_1_ and *m*_2_, we combined the corresponding populations resulting in a data set with 718 × 2 subejcts. In this case, the chance accuracy was equal to 50%. In the second pipeline, we employed an eight-class classificaiton task over the entire data set (excluding REST2). REST2 was excluded to eliminate the reduncdancy of resting-state session and balance the probability space evenly across sessions (chance accuracy = 12.5%). A predictive model was developed using gradient boosting machine [GBM (40)] with 100 estimators (or decision trees) and 0.05 learning rate. The entire population was randomly divided into two equal-size folds (each with 359 subjects): the training set and the testing set. The exemplar-search algorithm (the first part of the parcellation pipeline) was employed on the training set and the resulting exemplars were used to parcellate the entire population. Next, a GBM was trained on the training set using NNAs as features and the functional state as the output. The model was then tested to predict the functional state of the unseen subjects in the testing set. Both predictive pipelines were employed for K = 2 – 50. Note that here the exemplar identification was restricted to the training set. This was to assure that the training and the testing sets are independent throughout the entire pipeline. However, the same results were achieved using the initial parcellation schemes and employing a 10-fold GBM on the entire data set (Figure S4). The predictive pipeline was implemented using Python’s Scikit-learn library (82).

### State-decoding predictive features

Using GBM as our classifier facilitates retrieving the importance of features in the predictive analysis. Importance is explicitly calculated as the number of times that each feature was used to make decisions that improved the performance measure. This is weighted by the number of observations within each decision tree, and then averaged across all of the trees within the model. Since our GBM model was trained and tested over 268 features indicating the network assignment of each node, the importance of each node was assessed by the corresponding importance attribute. The importance of nodes within each entropy class was further assessed and compared using two-tailed t-test Bonferroni corrected for multiple comparisons.

### Cross-subject and cross-state entropies

From the NNA histograms derived earlier, we quantified the changes in the NNAs. To measure how frequently a node changes its memberships to a network, we used entropy, a well-established measure from the information theory literature (39). We defined and calculated two measures of entropy: cross-state and cross-subject entropy. Cross-state entropy was defined as the entropy of each row of matrix *P*_100×9_ summed up over all iterations:

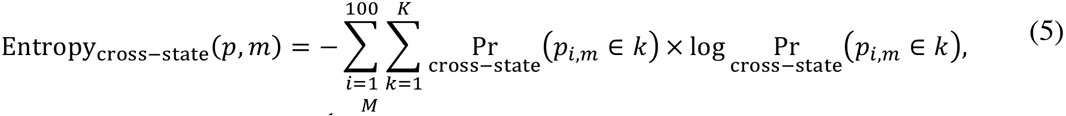

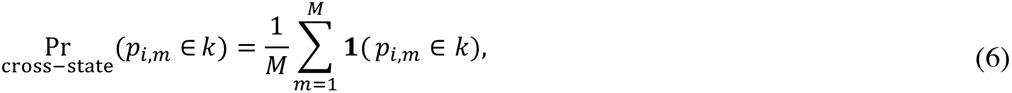

where **1** represents the indicator function, i.e.:

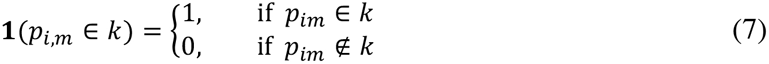

Here, 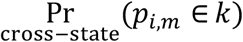 indicates the probability of node *p* in iteration *i* and state *m* being assigned to network *k*. This probability is calculated as the normalized NNA histogram across all states (Eq. 6).

Cross-subject entropy is defined as the entropy of each column of matrix *P*_100×9_ summed up over all states:

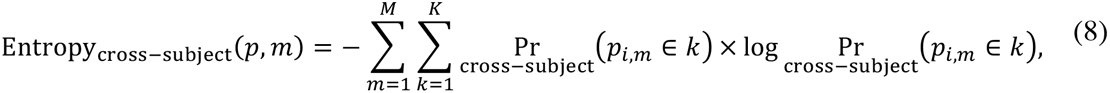

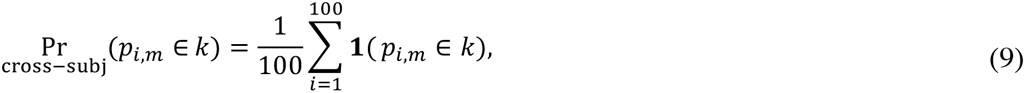

Here, 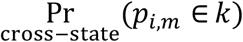 indicates the probability of node *p* in iteration *i* and state *m* being assigned to network *k*. This probability is calculated as the normalized NNA histogram across all iterations (Eq. 9). We further scaled the entropies to the range (0,100). Thus, every node is represented by two measures: cross-state and cross-subject entropies (Figure S5). These two measures were further exploited to group the nodes into three entropy classes: 1) *steady* nodes, those with low cross-state and cross-subject entropies, 2) *flexible* nodes, those with relatively high cross-state entropies, but low cross-subject entropies, and 3) *transient* nodes, regions with high cross-state and cross-subject entropies. Defining these entropy classes required a threshold for low and high entropies. We defined the steady nodes as regions with cross-state and cross-subject entropies equal to 0. These regions do not change their network assignments across different states or cohorts of subjects. In order to define the other two entropy classes, we took the mean of the cross-subject entropy among the rest of the non-steady nodes as the threshold (*τ*_mean_= 29.47). Thus, regions with cross-state entropies higher than 0 and cross-subject entropies lower than *τ*_mean_ were identified as flexible, and regions with cross-state entropies higher than 0 and cross-subject entropies higher than *τ*_mean_ were identified as transient. The more formal definition of the three entropy classes are as follows:

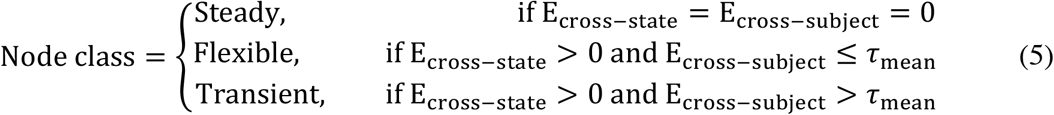

Where E stands for Entropy. These regions are further displayed on the brain with red indicating the steady nodes (n=72), yellow indicating the flexible nodes (n=115), and green representing the transient nodes (n=81).

### Localization of the node entropy classes in networks

We localized the entropy classes on the brain using two separate pipelines. In the first pipeline, we studied the distribution of different entropy classes within each network. For each system, the ratio of each of the three classes were displayed as a pie chart. In the second pipeline, we examined the contribution of the systems in each of the three classes by calculating the ratio of the nodes within every network assigned to each of the entropy classes. We reported the results using the network labels defined at REST1 as resting-state represents the intrinsic functional organization where these networks were originally developed (7, 9).

### Distribution of behavioral domains and paradigm classes across entropy classes

Behavioral domain (BD) and paradigm class (PC) profiles were defined by referencing the BrainMap Functional Database’s experimental meta-data [www.brainmap.org], which represents 62,038 x-y-z foci from 10,467 functional human brain imaging task-activation experiments representing 27,820 subjects. These task-activation experiments represent 55 BDs and 108 PCs (83, 84). BDs include categories and subcategories of mental processes isolated by the experimental contrasts. They comprise five main categories: cognition, action, perception, emotion, and interception. PCs include the experimental tasks isolated by an experimental contrast (see http://brainmap.org/scribe for more information on the BrainMap taxonomy). Profiles were computed within each node (or network) as a z-score of the forward inference likelihood that a particular BD or PC is reported within that node (or network) [*P(Activation | BD) or P(Activation | PC)*] normalized by the likelihood expected if they were uniformly distributed throughout the brain (85). A high z-score indicates a high specificity of a particular BD or PC for that node (or network). To create a profile for each node (or network), z-scores were thresholded at z>1.96 (associated with 95% confidence interval or p<0.05) and summed over all BD/PCs. Profiles were computed using MATLAB.

### Graph visualization of the functional connectivity reconfiguration across states

To provide a complementary analysis on how different networks interact with each other, we employed a graph visualization approach. While the delineation of functional networks represents the integration and segregation of different regions, functional connectivity analysis – using graph visualization – provides additional information on connections both within-and between-networks. This analysis further differentiates the notion of *community* and *connection,* as motivated by previous studies (33, 86). To this end, we constructed a population-level connectivity matrix for each state by taking the average of the individualized connectivity matrices (see Method 3.2). The average matrix was simplified by summarizing the edges to the top 10% and removing the within node connections (or loops) that were the artifact of correlation of vector by itself. The resulting matrix for each functional condition was visualized as a graph with the nodes colored according to their network assignments and the edges colored according to the entropy class of their two ending nodes. The homogenous connections – i.e. edges with the two ending nodes belonging to the same entropy class – are colored as red, for connections between two steady nodes, yellow for connections between two flexible nodes, and green for connections between two transient nodes. The heterogeneous connections – i.e. edges whose ending nodes belong to two different entropy classes – are colored using grayscale shades with dark gray indicating the connections between steady and flexible nodes, gray indicating the connections between steady and transient nodes, and light gray indicating the connections between flexible and transient nodes. The strength of these six classes of connections were assessed and compared using pair-wise two-tailed t-test, Bonferroni corrected for multiple comparisons. We further employed a test-retest analysis by repeating the entire graph visualization pipeline over two sets of 359 subjects derived by randomly splitting the entire population into two halves, and very similar results were qc. The graph visualization was employed using R package networkd3 [https://CRAN.R-project.org/package=networkD3].

## Reference

1. Brodmann K (1909) Vergleichende Lokalisationslehre der Grosshirnrinde in ihren Prinzipien dargestellt auf Grund des Zellenbaues (Barth).

2. Craddock RC, James GA, Holtzheimer PE, Hu XP, & Mayberg HS (2012) A whole brain fMRI atlas generated via spatially constrained spectral clustering. Human brain mapping 33(8):1914–1928.

3. Fan L, et al. (2016) The human brainnetome atlas: a new brain atlas based on connectional architecture. Cerebral cortex 26(8):3508–3526.

4. Glasser MF, et al. (2016) A multi-modal parcellation of human cerebral cortex. Nature 536(7615):171–178.

5. Shen X, Tokoglu F, Papademetris X, & Constable RT (2013) Groupwise whole-brain parcellation from resting-state fMRI data for network node identification. Neuroimage 82:403–415.

6. Finn ES, et al. (2015) Functional connectome fingerprinting: identifying individuals using patterns of brain connectivity. Nature neuroscience.

7. Power JD, et al. (2011) Functional network organization of the human brain. Neuron 72(4):665–678.

8. Smith SM, et al. (2009) Correspondence of the brain’s functional architecture during activation and rest. Proceedings of the National Academy of Sciences 106(31):13040–13045.

9. Thomas Yeo BT, et al. (2011) The organization of the human cerebral cortex estimated by intrinsic functional connectivity. Journal of Neurophysiology 106(3):1125–1165.

10. Meunier D, Lambiotte R, Fornito A, Ersche K, & Bullmore ET (2009) Hierarchical modularity in human brain functional networks. Frontiers in neuroinformatics 3:37.

11. Power JD, et al. (2011) Functional network organization of the human brain. Neuron 72(4):665–678.

12. Smith SM, et al. (2012) Temporally-independent functional modes of spontaneous brain activity. Proceedings of the National Academy of Sciences 109(8):3131–3136.

13. Dosenbach NU, et al. (2007) Distinct brain networks for adaptive and stable task control in humans. Proceedings of the National Academy of Sciences 104(26):11073–11078.

14. Laird AR, et al. (2011) Behavioral interpretations of intrinsic connectivity networks. Journal of cognitive neuroscience 23(12):4022–4037.

15. Grayson DS, et al. (2014) Structural and functional rich club organization of the brain in children and adults. PloS one 9(2):e88297.

16. Greicius MD, Srivastava G, Reiss AL, & Menon V (2004) Default-mode network activity distinguishes Alzheimer’s disease from healthy aging: evidence from functional MRI. Proceedings of the National Academy of Sciences of the United States of America 101(13):4637–4642.

17. Stern ER, Fitzgerald KD, Welsh RC, Abelson JL, & Taylor SF (2012) Resting-state functional connectivity between fronto-parietal and default mode networks in obsessive-compulsive disorder. PloS one 7(5):e36356.

18. van Eimeren T, Monchi O, Ballanger B, & Strafella AP (2009) Dysfunction of the default mode network in Parkinson disease: a functional magnetic resonance imaging study. Archives of neurology 66(7):877–883.

19. Mennes M, Kelly C, Colcombe S, Castellanos FX, & Milham MP (2012) The extrinsic and intrinsic functional architectures of the human brain are not equivalent. Cerebral cortex 23(1):223–229.

20. Krienen FM, Yeo BT, & Buckner RL (2014) Reconfigurable task-dependent functional coupling modes cluster around a core functional architecture. Phil. Trans. R. Soc. B 369(1653):20130526.

21. Cole MW, Bassett DS, Power JD, Braver TS, & Petersen SE (2014) Intrinsic and task-evoked network architectures of the human brain. Neuron 83(1):238–251.

22. Cole MW, Ito T, Bassett DS, & Schultz DH (2016) Activity flow over resting-state networks shapes cognitive task activations. Nature neuroscience 19(12):1718.

23. Telesford QK, et al. (2016) Detection of functional brain network reconfiguration during task-driven cognitive states. NeuroImage 142:198–210.

24. Cole MW, Bagic A, Kass R, & Schneider W (2010) Prefrontal dynamics underlying rapid instructed task learning reverse with practice. The Journal of neuroscience 30(42):14245–14254.

25. Cole MW, et al. (2013) Multi-task connectivity reveals flexible hubs for adaptive task control. Nature neuroscience 16(9):1348–1355.

26. Davison EN, et al. (2015) Brain network adaptability across task states. PLoS Comput Biol 11(1):e1004029.

27. Di X, Gohel S, Kim EH, & Biswal BB (2013) Task vs. rest—different network configurations between the coactivation and the resting-state brain networks.

28. Shine JM, et al. (2016) The dynamics of functional brain networks: integrated network states during cognitive task performance. Neuron 92(2):544–554.

29. Gratton C, et al. (2018) Functional brain networks are dominated by stable group and individual factors, not cognitive or daily variation. Neuron 98(2):439–452. e435.

30. Cohen JR & D’Esposito M (2016) The segregation and integration of distinct brain networks and their relationship to cognition. Journal of Neuroscience 36(48):12083–12094.

31. Mohr H, et al. (2016) Integration and segregation of large-scale brain networks during short-term task automatization. Nature communications 7:13217.

32. Bassett DS, et al. (2011) Dynamic reconfiguration of human brain networks during learning. Proceedings of the National Academy of Sciences 108(18):7641–7646.

33. Mattar MG, Cole MW, Thompson-Schill SL, & Bassett DS (2015) A functional cartography of cognitive systems. PLoS computational biology 11(12):e1004533.

34. Gordon EM, Laumann TO, Adeyemo B, & Petersen SE (2017) Individual variability of the system-level organization of the human brain. Cerebral Cortex 27(1):386–399.

35. Gordon EM, et al. (2017) Precision functional mapping of individual human brains. Neuron 95(4):791–807. e797.

36. Salehi M, Karbasi A, Shen X, Scheinost D, & Constable RT (2017) An exemplar-based approach to individualized parcellation reveals the need for sex specific functional networks. Neuroimage, Submitted 142.

37. Finn ES, et al. (2015) Functional connectome fingerprinting: identifying individuals using patterns of brain connectivity. Nature neuroscience 18(11):1664–1671.

38. Hamming R (1950) Error detecting and error correcting codes. Bell System Technical Journal 29:147–160.

39. Borda M (2011) Fundamentals in information theory and coding (Springer Science & Business Media).

40. Friedman JH (2001) Greedy function approximation: a gradient boosting machine. Annals of statistics:1189–1232.

41. Fox PT & Lancaster JL (2002) Mapping context and content: the BrainMap model. Nature Reviews Neuroscience 3(4):319.

42. Ito T, et al. (2017) Cognitive task information is transferred between brain regions via resting-state network topology. bioRxiv:101782.

43. Noble S, et al. (2017) Influences on the test–retest reliability of functional connectivity MRI and its relationship with behavioral utility. Cerebral Cortex 27(11):5415–5429.

44. Bijsterbosch JD, et al. (2018) The relationship between spatial configuration and functional connectivity of brain regions. Elife 7.

45. Zanto TP & Gazzaley A (2013) Fronto-parietal network: flexible hub of cognitive control. Trends in cognitive sciences 17(12):602–603.

46. Mueller S, et al. (2013) Individual variability in functional connectivity architecture of the human brain. Neuron 77(3):586–595.

47. Andrews-Hanna JR, Smallwood J, & Spreng RN (2014) The default network and self-generated thought: component processes, dynamic control, and clinical relevance. Annals of the New York Academy of Sciences 1316(1):29–52.

48. Zhang J, et al. (2016) Neural, electrophysiological and anatomical basis of brain-network variability and its characteristic changes in mental disorders. Brain 139(8):2307–2321.

49. Van Den Heuvel MP & Sporns O (2011) Rich-club organization of the human connectome. Journal of Neuroscience 31(44):15775–15786.

50. Park H-J & Friston K (2013) Structural and functional brain networks: from connections to cognition. Science 342(6158):1238411.

51. Bassett DS, et al. (2013) Task-based core-periphery organization of human brain dynamics. PLoS computational biology 9(9):e1003171.

52. Fedorenko E & Thompson-Schill SL (2014) Reworking the language network. Trends in cognitive sciences 18(3):120–126.

53. Crossley NA, et al. (2013) Cognitive relevance of the community structure of the human brain functional coactivation network. Proceedings of the National Academy of Sciences 110(28):11583–11588.

54. Anderson ML, Kinnison J, & Pessoa L (2013) Describing functional diversity of brain regions and brain networks. Neuroimage 73:50–58.

55. Power JD, Schlaggar BL, Lessov-Schlaggar CN, & Petersen SE (2013) Evidence for hubs in human functional brain networks. Neuron 79(4):798–813.

56. Li Y, et al. (2009) Brain anatomical network and intelligence. PLoS computational biology 5(5):e1000395.

57. Cohen JD, Lee RF, Norman KA, & Turk-Browne NB (2015) Closed-loop training of attention with real-time brain imaging. Nature neuroscience 18(3):470–475.

58. Rosenberg MD, et al. (2016) A neuromarker of sustained attention from whole-brain functional connectivity. Nature neuroscience 19(1):165–171.

59. Adelstein JS, et al. (2011) Personality is reflected in the brain’s intrinsic functional architecture. PloS one 6(11):e27633.

60. Woolgar A, Thompson R, Bor D, & Duncan J (2011) Multi-voxel coding of stimuli, rules, and responses in human frontoparietal cortex. Neuroimage 56(2):744–752.

61. Cole MW, Etzel JA, Zacks JM, Schneider W, & Braver TS (2011) Rapid transfer of abstract rules to novel contexts in human lateral prefrontal cortex. Frontiers in human neuroscience 5.

62. Heinzle J, Wenzel MA, & Haynes J-D (2012) Visuomotor functional network topology predicts upcoming tasks. Journal of Neuroscience 32(29):9960–9968.

63. Haxby JV, Connolly AC, & Guntupalli JS (2014) Decoding neural representational spaces using multivariate pattern analysis. Annual review of neuroscience 37:435–456.

64. Poldrack RA, Halchenko YO, & Hanson SJ (2009) Decoding the large-scale structure of brain function by classifying mental states across individuals. Psychological Science 20(11):1364–1372.

65. Perez Velazquez JL (2017) Dynamiceuticals: The Next Stage in Personalized Medicine. Frontiers in neuroscience 11:329.

66. Satterthwaite TD & Davatzikos C (2015) Towards an individualized delineation of functional neuroanatomy. Neuron 87(3):471–473.

67. Bzdok D & Meyer-Lindenberg A (2017) Machine learning for precision psychiatry: Opportunites and challenges. Biological Psychiatry: Cognitive Neuroscience and Neuroimaging.

68. Vu M-AT, et al. (2018) A Shared Vision for Machine Learning in Neuroscience. Journal of Neuroscience 38(7):1601–1607.

69. Emmert K, et al. (2016) Meta-analysis of real-time fMRI neurofeedback studies using individual participant data: How is brain regulation mediated? Neuroimage 124:806–812.

70. Hartwell KJ, et al. (2016) Individualized real-time fMRI neurofeedback to attenuate craving in nicotine-dependent smokers. Journal of psychiatry & neuroscience: JPN 41(1):48.

71. Koizumi A, et al. (2017) Fear reduction without fear through reinforcement of neural activity that bypasses conscious exposure. Nature human behaviour 1(1):0006.

72. Fitzgerald PB (2011) The emerging use of brain stimulation treatments for psychiatric disorders. Australian & New Zealand Journal of Psychiatry 45(11):923–938.

73. Plow E, et al. (2016) Models to tailor brain stimulation therapies in stroke. Neural Plasticity 2016.

74. Van Essen DC, et al. (2013) The WU-Minn human connectome project: an overview. Neuroimage 80:62–79.

75. Uğurbil K, et al. (2013) Pushing spatial and temporal resolution for functional and diffusion MRI in the Human Connectome Project. Neuroimage 80:80–104.

76. Rosenberg M, Noonan S, DeGutis J, & Esterman M (2013) Sustaining visual attention in the face of distraction: A novel gradual-onset continuous performance task. Attention, Perception, & Psychophysics 75(3):426–439.

77. Rosenberg MD, Hsu W-T, Scheinost D, Constable RT, & Chun MM (2017) Connectome-based models predict separable components of attention in novel individuals. The Journal of Cognitive Neuroscience, Submitted.

78. Shen X, et al. (2017) Using connectome-based predictive modeling to predict individual behavior from brain connectivity. nature protocols 12(3):506–518.

79. Smith SM, et al. (2013) Resting-state fMRI in the human connectome project. Neuroimage 80:144–168.

80. Glasser MF, et al. (2013) The minimal preprocessing pipelines for the Human Connectome Project. Neuroimage 80:105–124.

81. Joshi A, et al. (2011) Unified framework for development, deployment and robust testing of neuroimaging algorithms. Neuroinformatics 9(1):69–84.

82. Pedregosa F, et al. (2011) Scikit-learn: Machine learning in Python. Journal of machine learning research 12(Oct):2825–2830.

83. Barron D & Fox P (2015) BrainMap Database as a Resource for Computational Modeling.

84. Fox PT, et al. (2005) BrainMap taxonomy of experimental design: description and evaluation. Human brain mapping 25(1):185–198.

85. Lancaster JL, et al. (2012) Automated regional behavioral analysis for human brain images. Frontiers in neuroinformatics 6:23.

86. Bassett DS, Yang M, Wymbs NF, & Grafton ST (2015) Learning-induced autonomy of sensorimotor systems. Nature neuroscience 18(5):744–751.

